# ALS-linked SOD1 Mutants Enhance Outgrowth, Branching, and the Formation of Actin-Based Structures in Adult Motor Neurons

**DOI:** 10.1101/303271

**Authors:** Zachary Osking, Jacob I. Ayers, Ryan Hildebrandt, Kristen Skruber, Hilda Brown, Daniel Ryu, Amanda R. Eukovich, Todd E. Golde, David R. Borchelt, Tracy-Ann Read, Eric A. Vitriol

## Abstract

Amyotrophic lateral sclerosis (ALS) is a progressive, fatal neurodegenerative disease characterized by motor neuron cell death. However, not all motor neurons are equally susceptible. Most of what we know about the surviving motor neurons comes from gene expression profiling, less is known about their functional traits. We found that resistant motor neurons cultured from SOD1 ALS mouse models have enhanced axonal outgrowth and dendritic branching. They also have an increase in the number and size of actin-based structures like growth cones and filopodia. These phenotypes occur in cells cultured from presymptomatic mice and mutant SOD1 models that do not develop ALS, but not in embryonic motor neurons. Enhanced outgrowth and upregulation of filopodia can be induced in wild-type adult cells by expressing mutant SOD1. These results demonstrate that mutant SOD1 can enhance the regenerative capability of ALS resistant motor neurons. Capitalizing on this mechanism could lead to new therapeutic strategies.

## INTRODUCTION

Amyotrophic lateral sclerosis (ALS) is a fatal, adult-onset neurodegenerative disorder in which there is selective loss of motor neurons in the cerebral cortex, brainstem and spinal cord (Ince et al., 1998). Approximately 90% of ALS cases are sporadic with unknown etiology; the remaining 10% are inherited and known as familial ALS (fALS), of which over 20% have mutations in the gene encoding Cu/Zn superoxide dismutase 1 (SOD1) (Brown and Al-Chalabi, 2017). To date, over 155 different mutations have been identified in SOD1 either in isolated cases of ALS or more commonly in patients from families showing autosomal dominant patterns of inheritance (Andersen and Al-Chalabi, 2011; Pasinelli and Brown, 2006). ALS-linked SOD1 mutations are thought to induce a toxic gain-of-function in the protein, which becomes prone to misfolding and subsequent aggregation (Karch et al., 2009; Saccon et al., 2013). However, expression of mutant SOD1 can affect a number of cellular processes, causing ER distress, mitochondrial dysfunction, excitotoxicity, defects in axonal transport, and inhibition of the proteasome (Ilieva et al., 2009). Despite being the first gene identified with mutations that cause fALS (Rosen et al., 1993) and providing the basis of the first ALS animal model (Gurney, 1994), there is still no consensus about how mutant SOD1 specifically alters motor neuron physiology.

While most studies have focused on the cellular mechanisms and genes that induce motor neuron death in ALS, less is known about the neurons that do survive, including their ability to resist stress-induced cell death and to compensate for dying motor neurons. Not all motor neurons are equally susceptible to cell death during ALS disease progression. ALS mostly targets motor neurons required for voluntary movement, whereas motor neurons of the autonomic system are less sensitive (Piccione et al., 2015). There is also a gradient of vulnerability among spinal motor neurons, where faster motor units become affected before slower muscle types (Pun et al., 2006). Motor neurons that are less ALS-susceptible can compensate for the cells that initially die by establishing new connections with the motor endplate, although many of these will eventually succumb to the disease (Schaefer et al., 2005). This selective neuronal vulnerability is present in both sporadic ALS and familial ALS and is also recapitulated in rodent models, such as the SOD1^G93A^ mouse (Gurney, 1994; Nimchinsky et al., 2000).

Most of our current knowledge about surviving spinal motor neurons in ALS mouse models has largely been generated by gene expression profiling tissue and cells (Bandyopadhyay et al., 2013; Brockington et al., 2013; de Oliveira et al., 2013; Ferraiuolo et al., 2007; Lobsiger et al., 2007; Saxena et al., 2009). However, these studies provide just a single snapshot of the motor neuron’s biology and only allow for inferences to be made about how changes in gene expression alter motor neuron physiology, allow them to resist degeneration, or compensate for dying neurons by forming new motor endplate attachments. In the current study, we sought to functionally characterize ALS-resistant motor neurons by culturing them *in vitro*, where we would be able to directly assess dynamic cellular properties such as outgrowth, branching, and regulation of the cytoskeleton.

## RESULTS

### Axon outgrowth and branching are increased in adult motor neurons from symptomatic SOD1-ALS mice

To functionally characterize ALS-resistant motor neurons, we isolated them from adult mice expressing human SOD1^G93A^ at low copy number (referred to as G93A-DL) (Acevedo-Arozena et al., 2011). This model expresses between 6 and 8 copies of the human SOD1^G93A^ transgene, resulting in onset of ALS symptoms around 9 months of age. We corroborated these findings with the more extensively studied SOD1^G93A^ high copy number mouse model (referred to as G93A), which expresses SOD1^G93A^ at around 3 fold that of the G93A-DL model. These mice develop symptoms more rapidly, with hind-limb paralysis seen as early as 5 months of age (Gurney, 1994). Using a well-characterized protocol for the high yield extraction of spinal motor neurons from adult mice (Beaudet et al., 2015), we established cultures of adult motor neurons from mutant SOD1 and non-transgenic mice (referred to as NTg). We then performed a large-scale quantitative analysis of these cells’ ability to extend new processes. Since the isolation protocol severs all established neuronal projections, this assay is a direct measure of *in vitro* neurite regeneration.

Motor neurons from G93A-DL mice displayed significantly increased outgrowth in comparison to age- and sex-matched NTg controls, both in axon length (~55% longer) and in overall neurite branching complexity (~three times as many intersections 60 μm from the soma center) (Figure 1b-d). Motor neurons isolated from late-stage G93A mice also demonstrated increased neurite branching and axonal outgrowth relative to NTg mice (Figure 1c,d). In contrast, motor neurons from adult mice overexpressing wild-type SOD1 exhibited a slight decrease in outgrowth and branching (Figure 1c,d). This was an important control since the SOD1^G93A^ mutant maintains its enzymatic ability to remove superoxide radicals (Nishida et al., 1994). The reduction in axon extension and branching in motor neurons overexpressing wild-type SOD1 is consistent with previous findings where ROS depletion results in negative effects on neurite outgrowth (Munnamalai and Suter, 2009). Thus, the enhanced regeneration seen in late stage motor neurons is specific to the SOD1^G93A^ ALS mice and occurs with high and low expression levels of the mutant gene.

**Figure 1.**
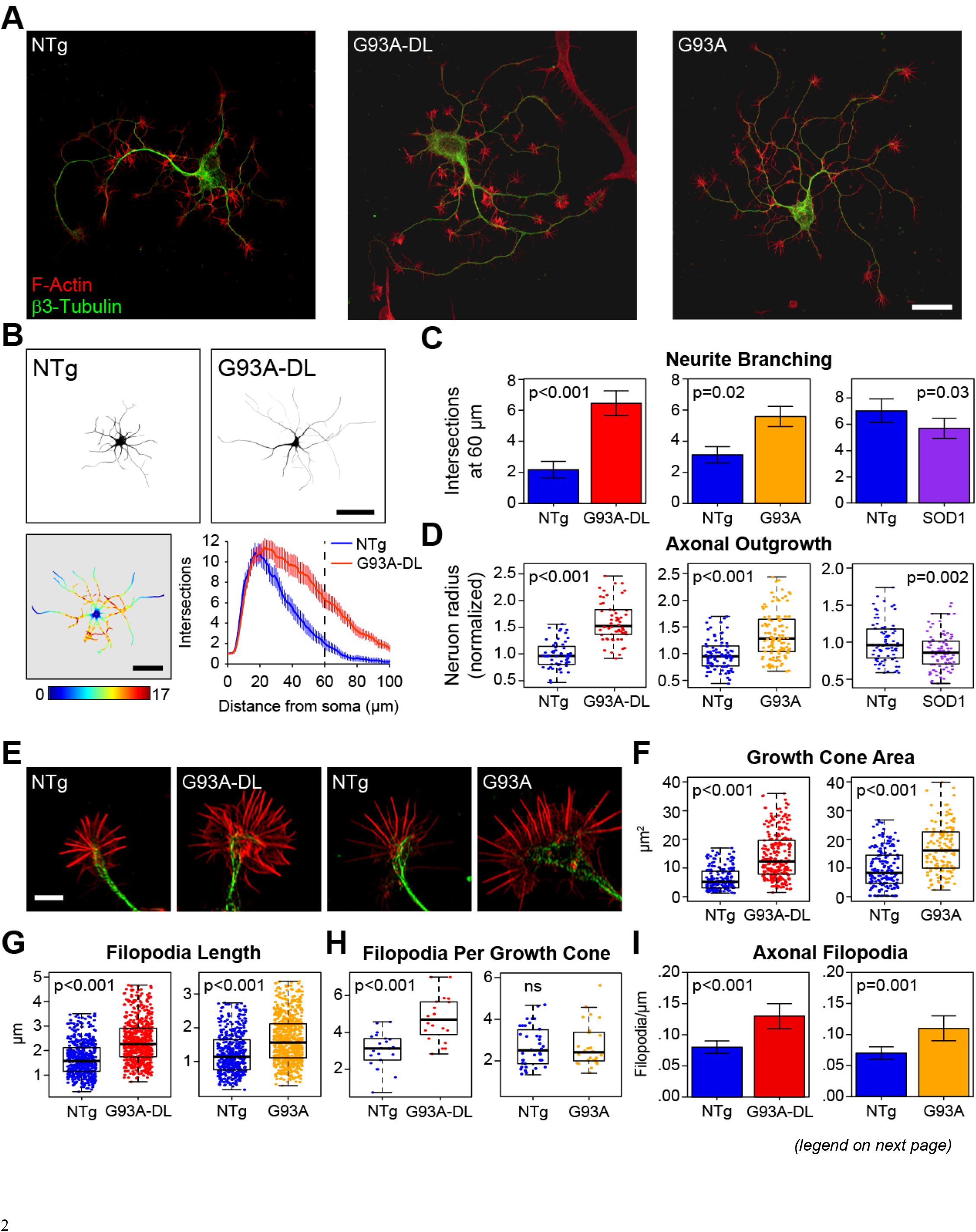
G93A and G93A-DL motor neurons from ALS symptomatic mice exhibit an enhanced outgrowth phenotype. (a) Epifluorescent images of representative motor neurons harvested from non-transgenic (NTg), low copy number SOD1^G93A^ overexpressing (G93A-DL), and SOD1^G93A^ overexpressing (G93A) mice. Motor neurons from both strains of SOD1^G93A^ mice were harvested at their respective symptomatic stages (approximately 9 months for G93A-DL and approximately 6 months for G93A). Scale bar is 30 μm. (b) Representative images of motor neurons harvested from NTg and ALS symptomatic G93A-DL mice. Pseudocolor image of G93A-DL motor neuron demonstrates the number of intersections measured at progressive distances from the soma. Scale bar is 50 μm. The line graph represents the average number of intersections measured at a given distance from the soma for all NTg or G93A-DL neurons. (c) Box-and-whisker plots of the axonal outgrowth (measured as the length of the longest neurite branch) of motor neurons cultured from G93A-DL, G93A mice and mice overexpressing SOD1^WT^ (SOD1) relative to their age-matched non-transgenic (NTg) controls. (d) Branching of motor neurons was quantified by measuring the average number of intersections found 60 μm from the center of the soma (noted as dotted line in (c)). Bar graphs depict motor neuron arborization seen in G93A-DL, G93A, and SOD1 motor neurons compared to their respective NTg controls For (c) and (d): G93A-DL data set: n=56 cells from 3 NTg mice and n=62 cells from 3 G93A-DL mice; G93A data set: n=88 cells from 4 NTg mice and n=120 cells from 5 G93A mice; SOD1 data set: n=82 cells from 2 NTg mice and n=89 cells from 3 SOD1 mice. (e) Representative images of G93A-DL and G93A growth cones of motor neurons harvested from symptomatic mice with NTg age-matched controls. Scale bar is 5 μm. (f-i) Plots depicting growth cone area, average filopodia length filopodia per growth cone, and axonal filopodia demonstrated in G93A and G93A-DL motor neurons from symptomatic mice relative to their NTg controls. For (f-g) n=25 for NTg, and n=25 for G93A-DL, harvested from two mice for each for G93A-DL data se and n=38 cells for NTg and n=28 cells for G93A, harvested from 2 mice each for G93A data set. For (i) n=46 cells from 4 mice for NTg and n=49 cells from 4 mice for G93A-DL, and n=30 cells from 2 mice for both NTg and G93A. Box-and-whisker plots denote the 95th (top whisker), 75th (top edge of box), 25th (bottom edge of box), and 5th (bottom whisker) percentiles, and the median (bold line in box). Each point on the graph represents an individual cell measurement, scattered with random noise in the x direction to enable visualization of all points for the given data set. Error bars correspond to the 95% confidence interval for each data set. p-values were obtained from a two-tailed student’s T-test.

### Actin-based structures are increased in adult motor neurons from symptomatic SOD1-ALS mice

Growth cones are the motile organelles found at the tip of axonal and dendritic projections and play a pivotal role in outgrowth and pathfinding. The peripheral region of the growth cone contains actin-based lamellipodia and filopodia, two types of membrane protrusions that function in growth cone movement and environment sensing (Vitriol and Zheng, 2012). We observed a marked increase in the size of growth cones and filopodia in spinal motor neurons isolated from symptomatic G93A-DL and G93A mice. G93A-DL growth cones were on average greater than twice the size of those from NTg controls, G93A growth cones exhibited a similar increase in size (Figure 1f). Growth cone filopodia from both ALS mouse lines were also significantly longer than NTg controls, with G93A-DL cells exhibiting the largest size difference (Figure 1g). G93A-DL growth cones also contained more filopodia (Figure 1h). Axonal filopodia are actin-based structures extending off of the main axon terminal that serve as precursors for collateral branches, which are involved in building complex neural circuits (Gallo, 2013). In ALS, the formation of new collateral branches occurs in the resistant motor units as they attempt to expand their synaptic connections to compensate for early denervation events (Clark et al., 2016; Schaefer et al., 2005). In motor neurons isolated from both G93A-DL and G93A mice, we observed a marked increase in axonal filopodia density relative to NTg controls (Figure 1i). Thus, the surviving motor neurons isolated from symptomatic ALS mice exhibit an upregulation of multiple actin-based structures associated with outgrowth and regeneration.

### Enhanced regeneration of mutant SOD1 motor neurons only occurs in adult cells and is independent of ALS onset

Our results were surprising since previous studies have shown that the expression of G93A is either inhibitory or has no effect on outgrowth and regeneration in motor neurons. However, these studies were conducted using embryonic cells (Nagai et al., 2007) or iPSC-derived motor neurons (Isobe et al., 2015; Karumbayaram et al., 2009), which more closely resemble the embryonic state (Ho et al., 2016), and may respond differently to the mutant SOD1 expression. When we cultured motor neurons from G93A-DL and NTg pups at E14, there was no significant difference observed in outgrowth or branching after 3 days *in vitro* (DIV) (Figure 2a,b). We then cultured motor neurons from adult G93A-DL mice at different time points prior to the onset of ALS symptoms (1, 2, and 6 months of age). Increased axonal outgrowth and neurite branching relative to NTg controls were observed the 2 and 6 months timepoints, with a more significant difference at 6 months (Figure 2d). This data reveals a trend where regeneration is enhanced relative to NTg controls as the mice age and become closer to developing ALS. Though, if the actual size of the processes are plotted instead of their relative size (to NTG controls), G93A-DL motor neurons maintain the same level of outgrowth (axon is ~120 μm) throughout their lifespan, while NTg motor neurons actually become progressively smaller. The same trend exists for branching (Figure 2d). This could be interpreted as G93A-DL motor neurons having a preserved, rather than an enhanced, ability to regenerate.

**Figure 2.**
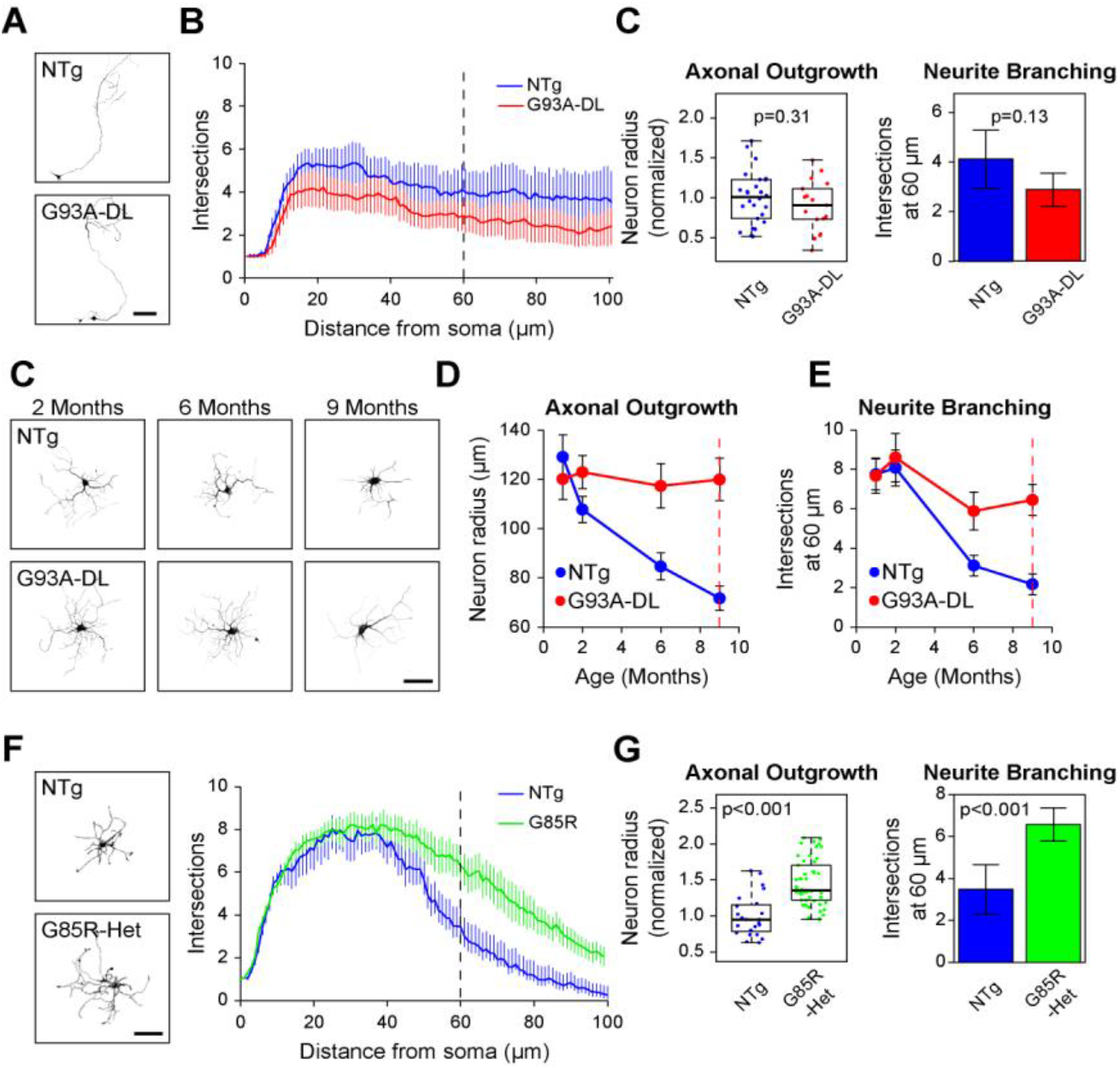
Increased outgrowth can be observed in mutant SOD1 motor neurons independently of the onset of ALS symptoms. (a) Representative images and Sholl graph of E14 motor neurons harvested from NTg and G93A-DL mice. Scale bar represents 100 μm. (b) Box-and-whisker plot and bar graph showing axonal outgrowth (neuron radius) and neurite branching (number of intersections found 60 μm from the center of the soma (noted as dotted line in (a)) measurements for embryonic NTg and G93A-DL motor neurons. (c) Representative images of adult motor neurons harvested at 2, 6, and 9 months from NTg and G93A-DL mice. Scale bar represents 50 μm. (d) Line graphs depicting axonal outgrowth and neurite branching as a function of age. For the data set taken at 2 months n=95 for NTg and n=80 for G93A-DL, harvested from 3 mice each. Data representing 6 months was taken from 3 mice each for NTg and G93A-DL, with n=88 and n=80 respectively. Data representing 9 months (symptomatic stage, indicated with dotted line) was taken from 3 NTG mice and 3 G93A-DL mice, with n=56 and n=62 respectively. (e) Representative images and Sholl graph of motor neurons harvested from NTg and G85R mice at 5 months of age. Scale bar represents 50 μm. (f) Box-and-whisker plot and bar graph showing axonal outgrowth (neuron radius) and neurite branching (number of intersections found 60 μm from the center of the soma (noted as dotted line in (e)) measurements for G85R and NTg motor neurons. n=23 from 2 mice for NTg and n=51 from 4 mice for G85R.

To verify that enhanced outgrowth of adult motor neurons from mutant SOD1 mouse models occurs independently of developing ALS, we isolated cells from heterozygous mice overexpressing YFP-SOD1G85R. YFP-SOD1G85R homozygous knock-in mice get ALS, while the heterozygous mice (referred to as G85R-het) do not develop symptoms (Bruijn et al., 1997; Wang et al., 2009a). Thus, the heterozygous model is a useful tool for studying the effects of mutant SOD1 overexpression independently of the effects of ALS progression. Motor neurons were isolated from mice at 5 months of age, the same age of ALS symptom presentation in the G93A model and an age in pre-symptomatic G93A-DL mice that still has substantial increases in outgrowth and branching relative to controls (Figure 1c,d, and 2d). G85R-het motor neurons exhibited an increase in axon length and neurite branching comparable to that seen in end-stage G93A mice (Figure 2e,f). Thus, expression of mutant SOD1 can enhance regeneration independently of ALS symptoms even in a model where there is no selection for surviving cells or stress from motor neuron death that signals the remaining neurons to reinnervate lost connections (Höke et al., 2006). This strongly suggests that it is the expression of mutant SOD1, not external factors caused by ALS, which increases outgrowth and regeneration of adult motor neurons.

### Expression of SOD1^G93A^ enhances outgrowth and branching of wild-type adult motor neurons

All of the experiments done so far had been performed with animal models where the cells had been expressing a mutant transgene for months. In fact, the enhanced outgrowth and branching phenotypes only becomes apparent after the mouse is two months old (Figure 2d). Therefore, it could be argued that enhanced regeneration is the result of an accumulated effect caused by long-term expression of the mutant gene, explaining the differences seen between adult (Figures 1-2) and embryonic motor neurons (Figure 2a,b). To determine if acute expression of SOD1^G93A^ was sufficient to increase outgrowth and branching in adult motor neurons, we cultured wild-type cells from 9-12 month old NTg mice and transduced them with adeno-associated virus (AAV) to express wild-type SOD1 (SOD1-YFP), SOD1^G93A^ (G93A-YFP), or a GFP control. Expression of G93A-YFP was sufficient to increase axonal outgrowth relative to NTg motor neurons (Figure 3b). While branching was not significantly increased, there was a significant positive correlation between G93A-YFP expression levels and both outgrowth parameters measured (Figure 3c). The difference between cells expressing G93A-YFP and SOD1-YFP was even more significant for outgrowth and branching (Figure 3b). SOD1-YFP positive cells did not have statistically significant differences in outgrowth and branching relative to GFP controls, but expression of SOD1-YFP was negatively correlated with outgrowth, mimicking the trend observed in the SOD1 overexpressing transgenic mouse models (Figure 1c,d). Thus, acute expression of mutant SOD1 was sufficient to increase regeneration of adult wild-type motor neurons.

**Figure 3.**
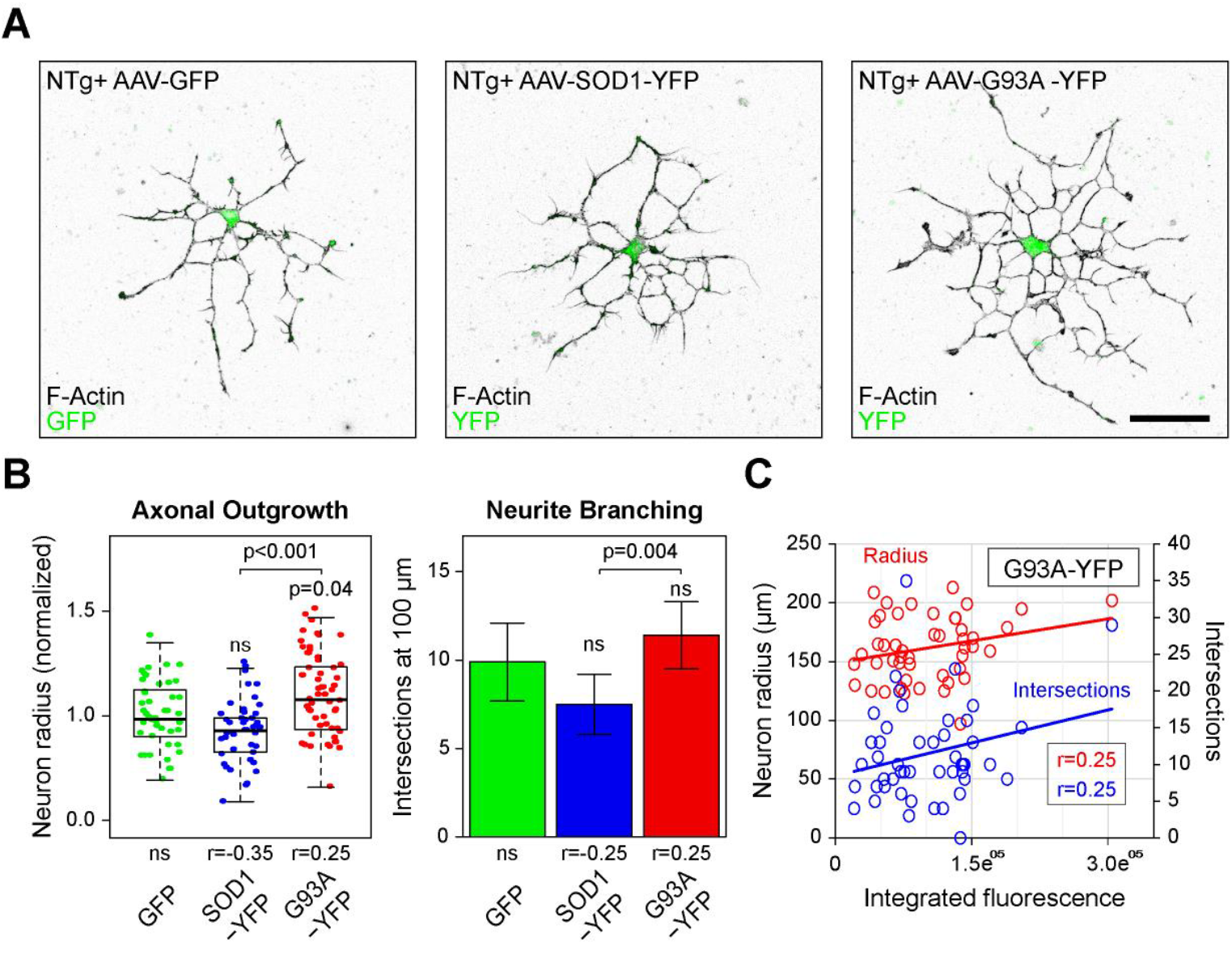
Acute expression of SOD1^G93A^ increases outgrowth in adult motor neurons. (a) Representative images of NTg motor neurons from 9-12 month old mice transduced with AAV to overexpress GFP, wild-type SOD1-YFP, or SOD1G93A-YFP (G93A-YFP). Scale bar is 50 μm. (b) Box-and-whisker plot depicting axonal outgrowth (neuron radius) and bar graph showing branching (number of intersections found 100 μm from the center of the soma) measurements of transduced NTg motor neurons. n=38 cells from 6 mice, n=37 cells from 6 mice, and n=48 cells from 6 mice for AAV-GFP, AAV-SOD1-YFP, or AAV-G93A-YFP, respectively. (c) Scatterplot depicting the relationship between the mean cellular fluorescence intensity of G93A-YFP axon outgrowth, and neurite branching. n=46, n=45, and n=57, for NTg motor neurons infected with AAV-GFP, AAV-SOD1-YFP, or AAV-G93A-YFP respectively, cells harvested from 9 mice (3 trials, 3 mice/trial). p-values were obtained ANOVA followed by Tukey’s post-hoc test. R-values represent Pearson’s correlation coefficient, r-values were only reported if the correlation was statistically significant (P<0.05).

### SOD1^G93A^ increases axonal filopodia and localizes to actin-based structures

Since actin-based structures were upregulated in motor neurons from G93A mouse models (Figure 1f-i), we wanted to determine if expression of SOD1^G93A^ was sufficient to increase them in wild-type motor neurons. Cells from NTg mice were infected with AAV expressing GFP, SOD1-YFP, and G93A-YFP and then axonal filopodia were measured. Quantification of growth cone parameters was not possible due to length of time the cells were cultured to achieve robust transgene expression (10 DIV compared to 3 DIV for our analysis of end stage G93A/G93A-DL motor neurons). Growth cones are most prevalent at 3 DIV. By 10 DIV few cells still had growth cones, but filopodia were still ubiquitously present on neurite projections, so we quantified axonal filopodia density. Both SOD1-YFP and G93A-YFP expressing cells had significantly higher axonal filopodia densities compared to GFP, with G93A-YFP expressing cells having the largest increase in filopodia (Figure 4a). G93A-YFP expression resulted in a 33% increase in filopodia density over SOD1-YFP and a 100% increase over GFP (Figure 4a). Further, we observed that G93A-YFP was localized to actin-based structures such as growth cones (the few that were present) and filopodia more than SOD1-YFP (Figure 4b).

**Figure 4.**
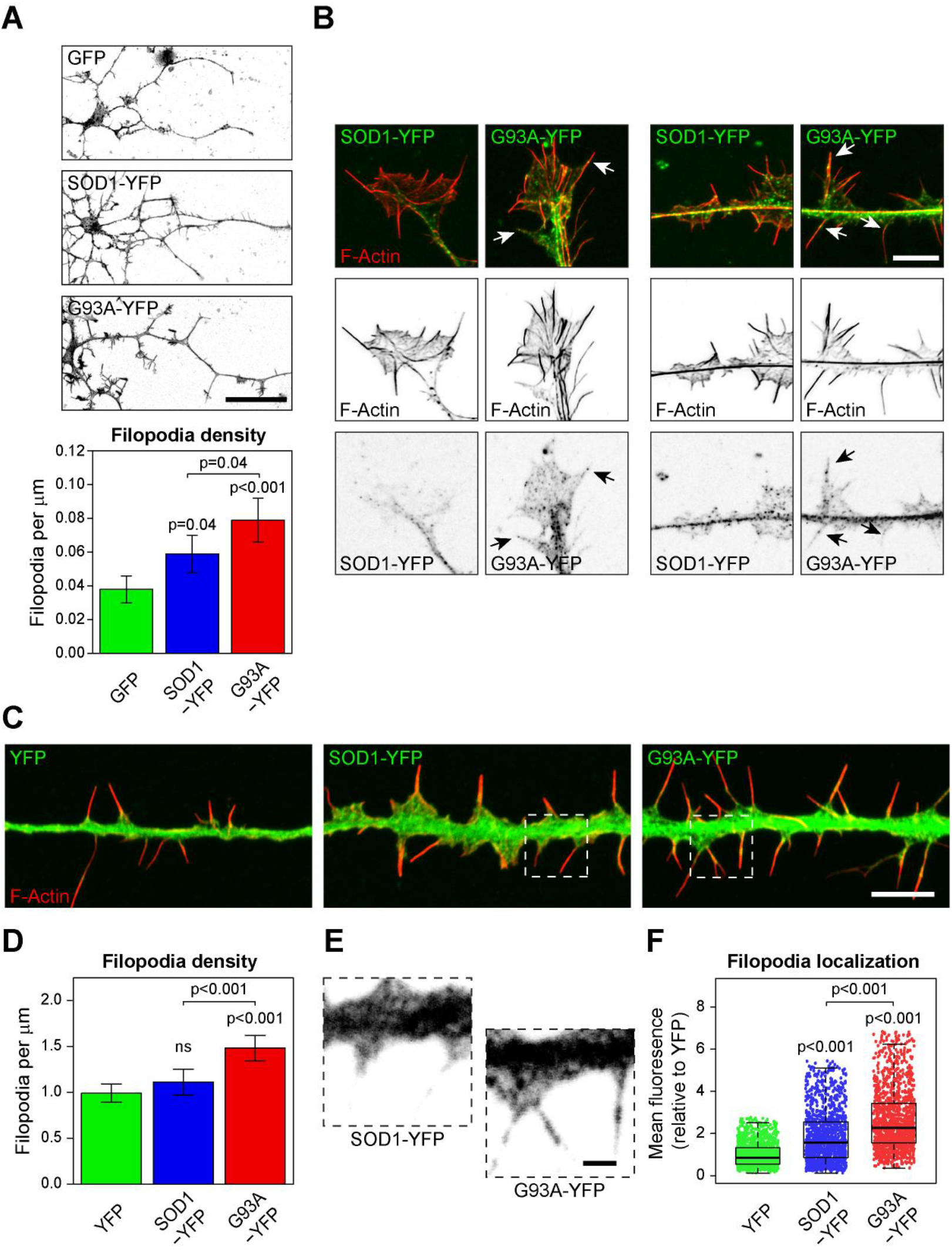
SOD1^G93A^ is preferentially integrated into filopodia and its overexpression results in increased filopodia density. **(a)** Representative images and quantification of axonal filopodia density (measured on the longest projection) from AAV-GFP, AAV-SOD1-YFP, or AAV-G93A-YFP transduced NTg motor neurons. n=38 cells from 6 mice, n=37 cells from 6 mice, and n=48 cells from 6 mice for AAV-GFP, AAV-SOD1-YFP, or AAV-G93A-YFP, respectively. Scale bar is 50 μm. **(b)** Confocal images of neurites from NTg adult motor neurons transduced with AAV-SOD1^WT^-YFP and AAV-SOD1^G93A^-YFP. Arrows indicate localization of G93A-YFP to filopodia. Scale bar is 5 μm. **(c)** Representative images of neurite-like projections in Cath-a-differentiated (CAD) cells that have been transfected with YFP, SOD1^WT^-YFP (SOD1-YFP), or SOD1^G93A^-YFP (G93A-YFP). Scale bar is 5 μm. **(d)** Bar graph measuring filopodia density in CAD cells transfected with YFP, SOD1-YFP, or G93A-YFP. n=71 cells for YFP, n=48 cells for SOD1-YFP, and n=72 for G93A-YFP, three separate transfections per condition. **(e)** Enlarged view of YFP fluorescence from dashed boxes from **(c)** showing the increase localization of SOD1^G93A^-YFP into filopodia compared to SOD1-YFP. Scale bar represents 1 μm. **(f)** Box-and-whisker plot quantifying YFP fluorescence intensity in filopodia in CAD cells. n=1359 filopodia from 38 cells for YFP, n=1367 filopodia from 40 cells for SOD1-YFP, and n=1524 from 39 cells for G93A-YFP, three separate transfections per condition.

To more robustly quantify how SOD1^G93A^ increases and localizes to filopodia, we overexpressed G93A-YFP, SOD1-YFP, or YFP in Cath.-a-differentiated (CAD) cells. CAD cells were differentiated for 18 hours prior to imaging so that they extended neurite-like structures which are highly similar to actual neural projections (Kapustina et al., 2016; Li et al., 2007). These neurite-like projections contain numerous filopodia. Filopodia were imaged using super-resolution confocal microscopy (Wilson, 2011) and then analyzed with the ImageJ plugin Filopodyan, which provides a platform for automated segmentation and analysis of filopodia (Urbančič et al., 2017). This enabled us to measure the YFP fluorescence of hundreds of individual filopodia per condition. We observed a significant increase in the localization of SOD1-YFP and G93A-YFP into filopodia over the YFP control, with G93A having the most robust filopodia localization (Figure 4e,f). Additionally, we observed an increase in filopodia density when G93A-YFP was expressed (Figure 4d) that was similar to experiments performed with adult motor neurons (Figure 4a). Thus, SOD1^G93A^ localizes to and increases filopodia in multiple cell types.

## DISCUSSION

This study demonstrates that SOD1^G93A^ expression can have pro-regenerative effects on adult motor neurons. Not only was enhanced neurite regeneration observed in motor neurons isolated from mutant SOD1 transgenic mice (Figure 2), but the expression of SOD1^G93A^ alone was sufficient to increase outgrowth in non-transgenic primary motor neurons (Figure 3). Further, SOD1^G93A^ localized to actin-based cellular structures and increased their size and number (Figure 4). Finally, an important take-home message from this study is the importance of working with the right cell type as mutant SOD1 expression had no effect on embryonic motor neuron regeneration (Figure 2a,b), but had a substantial effect on adult motor neuron outgrowth and branching (Figure 1-3). While it has been speculated that upregulation of regenerative/injury pathways is merely a compensatory response to mutant SOD1 induced toxicity (Lobsiger et al., 2007; Pun et al., 2006), our work suggests a novel gain-of-function for mutant SOD1, where it can help preserve motor neuron plasticity.

There are two mechanisms through which this could occur. First is a direct regulation of actin by mutant SOD1. While the relationship between mutant SOD1 and the actin cytoskeleton is not well characterized, it has been shown that SOD1^G93A^ directly interacts with actin in cultured cells (Takamiya et al., 2005) and spinal cord homogenates (Zetterstrom et al., 2011) by co-immunoprecipitation. Interestingly, mutant SOD1 has a significantly higher affinity for actin than the wild-type protein (Takamiya et al., 2005). That SOD1 is found in filopodia is also of interest. Since filopodia are extremely thin (~200 nm) extensions of the cellular membrane that are tightly packed with rearward-flowing actin filaments, localization there strongly indicates a specific interaction (Bird et al., 2017). It also might suggest that SOD1 preferentially binds or bundles linear arrays of filaments since the most prevalent structures we found to be upregulated by expression of mutant SOD1 were axonal filopodia. Axonal filipodia are precursor membrane protrusions to collateral branches (Gallo, 2013). New collateral branches form during the early stages of ALS, when the resistant neurons try to establish new synaptic connections to compensate for the denervation caused by the loss of the most susceptible neurons (Clark et al., 2016; Schaefer et al., 2005). Axonal filopodia must recruit microtubules in order to become collateral branches (Ketschek et al., 2015). Interestingly, in addition to actin, mutant SOD1 can also interact with the microtubule cytoskeleton (Kabuta et al., 2009). Thus, mutant SOD1 may have a dual role in the formation of new branches by increasing axonal filopodia (Figure 4a,d) and then helping microtubules to enter them. However, future studies will be required to determine if mutant SOD1 is directly involved in cytoskeletal regulation.

The second way that mutant SOD1 expression could enhance outgrowth and branching would be through upregulation of pro-regenerative signaling and cytoskeletal pathways. There are several published studies characterizing genetic changes in motor neurons from G93A mice at various stages of disease progression (D’Arrigo et al., 2010; de Oliveira et al., 2013; Ferraiuolo et al., 2007; Guipponi et al., 2010; Offen et al., 2009; Perrin et al., 2005; Saris et al., 2013; Yu et al., 2013). However, these studies have not reached consensus regarding the underlying genetics promoting ALS resistance, probably due to the variation in experimental design and tissue sampling. For example, using G93A mice, one study found a massive upregulation of genes involved in cell growth and/or maintenance in micro-dissected motor neurons from the lumbar spinal cord (Perrin et al., 2005), whereas another study found Wnt signaling to be significantly activated when homogenized whole spinal cord was used for RNA extraction (Yu et al., 2013). It has also been shown that upregulation of axonal guidance genes and actin cytoskeletal genes (including alpha- and beta-actin) occurs from the lumbar spinal cord of pre-symptomatic G93A mouse (de Oliveira et al., 2013). Thus, SOD1^G93A^ may prime adult motor neurons for outgrowth and regeneration through a positive activation of genes that regulate the actin cytoskeleton. However, it should be noted that many of the initially resistant motor neurons eventually succumb to ALS (Schaefer et al., 2005). The positive influence of mutant SOD1 expression on adult motor neuron regeneration most likely reflects an intermediate state occurring before cytotoxicity overwhelms the cells. However, instead of trying to completely eliminate it (Smith et al., 2006), capitalizing on the pro-regenerative effects mutant SOD1 while combating its toxic ones could be a useful strategy for future therapies.

## MATERIALS AND METHODS

### Mouse colony housing and breeding

All studies involving mice were approved by the Institutional Animal Care and Use Committee (IACUC) at the University of Florida in accordance with NIH guidelines. Adult mice were housed one to five per cage and maintained on *ad libitum* food and water, with a 12 h light/dark cycle. Transgenic mouse strains SOD1-G93A and SOD1-G93A-DL (JAX stock #004435 and #002299 respectively (Gurney, 1994) were purchased from Jackson laboratory (Bar Harbor, Maine), and bred by the Rodent Breeding Services offered by the Animal Care Services at the University of Florida. SOD1-G93A and SOD1-G93-DL colonies were maintained by breeding hemizygous mice either to wild type siblings, or to C57BL/6J inbred mice (Jax Stock # 000664). Additional transgenic mice strains used in this study were G85R-SOD1:YFP (Wang et al., 2009b) and WT-SOD1 (JAX stock #002297), both strains were generously supplied by Dr. Borchelt. The G85R-SOD1:YFP mice were maintained as heterozygotes on the FVB/NJ background and whereas the WT-SOD1 were maintained on a C57Bl/6J and C3H/HeJ hybrid background. In addition to the adult mice, we also used timed-pregnant C57BL/6J and SOD1-G93A-DL mice at gestational day 14, which were also generated by the Rodent Breeding Services offered by the Animal Care Services at the University of Florida.

Colony maintenance genotyping for all strains was performed as previously described (Gurney, 1994; Wang et al., 2003). Furthermore, to control for possible transgene copy loss due to meiotic rearrangement, breeders were regularly screened by RT-PCR as previously described (Henriques et al., 2010) and replaced with fresh founder stocks from Jackson laboratory (Bar Harbor, Maine) every 5 generations. In our colony SOD1-G93A and SOD1-G93-DL mice reached late disease stage at 150-180 days and 240-330 days of age respectively.

### Assessment of ALS disease progression

Mice were considered symptomatic if they displayed a 15% loss of bodyweight or showed signs of leg paralysis, whichever was reached first. In our hands, the majority of mice (~70%) were euthanized because of leg paralysis, and the rest due to decreased body weight. All mice were euthanized by CO_2_ inhalation following the guidelines provided by the University of Florida Animal Care Services (ACS) and approved by the Institutional Animal Care and Use Committee (IACUC). Late disease stage was defined by hind leg paralysis/weight loss.

### Study design

To control for sex differences in disease progression and phenotype of SOD1-G93A mice, symptomatic adult G93A and G93A-DL mice were always paired with non-transgenic (NTg) mice of the same sex and of similar age for each experiment, in most cases using littermates. Age and sex matching also allowed us to control for batch differences in the conditioned medium used to culture adult motor neurons as described below under “cell culture conditions”.

### Adult and embryonic mouse spinal cord isolation

Embryo spinal cords were obtained from timed pregnant G93A-DL and C57BL/6J mice at embryonic day 14 as previously described in detail (Beaudet et al., 2015). Once embryos were removed from the uterus, spinal cords were extracted under sterile conditions in a laminar flow hood with the aid of a dissecting microscope (SMZ800, NIKON INSTRUMENTS INC.) and small forceps and placed into cold Leibovitz’s L-15 medium (Life Technologies, Grand Island, NY) supplemented with 25 μg ml^−1^ penicillin-streptomycin (Life Technologies). The meninges and dorsal root ganglia (DRG) were peeled off and individual spinal cords were transferred into a 12 wells plate, identified and kept in cold L-15 medium on ice. Tails from each embryo were also harvested at this point for genotyping (G93A-DL mice).

Adult spinal cords were isolated by cutting the vertebrate column with scissors in front of the back legs and just below the medulla oblongata and flushed out of the spinal column using a syringe filled with cold supplemented DMEM/F12-medium with 18G needle (BD Biosciences). The DMEM/F12-medium used for this purpose consisted of DMEM/F12 in a 3:1 ratio supplemented with 36.54 mM NaHCO3 (Fisher Scientific), 0.18 mM L-adenine (Sigma), 312.5 μl L^−1^ 2N HCL (Fisher Scientific), 10% of fetal calf serum (Hyclone, GE Healthcare Life Sciences, South Logan, Utah) and 25 μg ml^−1^ of penicillin-streptomycin (Life Technologies). The adult spinal cords were transferred into cold DMEM/F12-medium.

### Motor neuron cell extraction and separation

Both embryonic and adult motor neurons were extracted using the method and reagents described in detail by Beaudet *et al*. (Beaudet et al., 2015) with a few modifications. Briefly, individual spinal cords were cut into small pieces and incubated for 30 min at 37°C in digestion buffer consisting of Dulbecco’s PBS (DPBS, Life Technologies, Grand Island, NY) containing 10 U/ml^−1^ papain (Worthington, Lakewood, NJ, USA), 200 μg/ml^−1^ L-cysteine (Sigma St. Louis, MO) and 250 U/ml^−1^ DNase (Sigma, St. Louis, MO). The digestion buffer was then removed and replaced with DPBS containing 8 mg/ml Ovomucoid trypsin inhibitor (Sigma), 8 mg/ml bovine serum albumin (BSA, Sigma), and 250 U/ml DNase. The tissue was then triturated using glass pipettes to obtain a single-cell suspension. This step was repeated trice before all cells were collected and filtered through a 40 μm cell strainer (BD Falcon) and centrifuged at 280 g for 10 min at 4 °C for motor neuron. Adult mixed motor neuron cultures were ready to plate after this step. Embryonic motor neuron pellets were enriched by resuspending in 6 ml of cold Leibovitz’s L-15 medium (Life Technologies) and laid over a 1.06 g ml−1 Nycoprep density solution (Axis-Shield, Dundee, Scotland) and spun at 900 g for 20 min at 4 °C without brake in a swinging bucket centrifuge (Eppendorf, Hauppauge, NY). Motor neurons were collected at the interface of the Nycoprep solution and poured in a new 50 ml collection tube which was then filled with cold L-15. Motor neuron cells were counted at this step. Motor neuron collecting tubes were centrifuged at 425 g for 10 min in a swinging bucket centrifuge at 4°C.

### Cell culture

Embryonic motor neuron pellets were gently resuspended at 200,000 cells/cm^2^ in freshly prepared Motor Neuron Growth Medium (MNGM), which is described in detail by Graber DJ *et al* (Graber and Harris, 2013). Briefly, the MNGM consists of Neurobasal A medium (NB-medium, Life Technologies) supplemented with 1X B-27 Serum-Free Supplement (Gibco/Life Technologies), 1X SATO supplement, 5 μg mL^−1^ Insulin (Gibco/Life Technologies), 1 mM Sodium pyruvate (Gibco/Life Technologies), 2 mM L-Glutamine (Gibco/Life Technologies), 40 ng mL^−1^ of 3,3,5-triiodo-L-thyronine sodium salt (T3; Sigma-Aldrich), 1 μg mL^−1^ Mouse laminin (Gibco/Life Technologies), 417 ng mL^−1^ Forskolin (Sigma-Aldrich), 5 μg mL^−1^ N-acetyl-L-cysteine (NAC, Sigma-Aldrich) and 1x Penicillin-streptomycin (Gibco/Life Technologies). After filter-sterilization using a 22 μm syringe filter, 10 ng mL^−1^ of each of the following growth factors was added to the medium: brain-derived neurotrophic factor (BDNF; Sigma-Aldrich), ciliary neurotrophic factor (CNTF; Peprotech, Rocky Hill, NJ) and glial-derived neurotrophic factor (GDNF; Peprotech). Embryonic motor neurons were either seeded on to Poly-D-lysine (PDL) coated 6cm tissue culture plates (10 μg mL^−1^ PDL, Sigma-Aldrich) to generate conditioned medium used for adult motor neuron cultures, or on to 1.5 cm glass coverslips pre-coated first with PDL (10 μg mL^−1^ for 1h at RT) then with Human Placental Laminin for 3 h at 37°C (1.67 μg mL^−1^ laminin in NB-Medium, Sigma-Aldrich). Embryonic motor neurons grown on coverslips were cultured for 3 days prior to fixation in 4% PFA and immunostaining for imaging and growth analysis.

Adult mixed motor neuron cultures were seeded onto PDL coated cover slips (10 μg mL^−1)^ and cultured in MNGM which had been pre-conditioned for 4 days by embryonic motor neurons isolated from NTg C57BL/6J mice. Given that adult motor neuron pellets contain considerable amount of debris when first plated, cells were not counted prior to seeding. After the cells were allowed to attach to the coverslips in a humidified 37°C incubator for 1 h, they were washed twice with warm NB-medium to remove debris and cultured in 1 ml of conditioned MNGM mixed 1:1 with freshly prepared MNGM. Adult motor neurons were cultured for 2 days prior to fixation and immunostaining for imaging and growth analysis. To confirm that these cultures were indeed enriched with motor neurons, cells were cultured for two days *in vitro* and immunostained for the motor neuron specific markers choline acetyltransferase (ChAT) or LIM-homeobox gene islet-1 (Isl1) (Supplemental Figure 1). Motor neurons where selected for analysis based on their expression of β3 tubulin, their large size, multi-polarity, and stellate cell shape.

Cath.-a-differentiated (CAD) cells (purchased from Sigma-Aldrich) cells were cultured in DMEM/F12 medium (Gibco) supplemented with 8% fetal calf serum, 1% L-Glutamine, and 1% penicillin-streptomycin. CAD cells were differentiated in the same medium without serum. They were imaged in DMEM/F12 medium without phenol red (Gibco) supplemented with 15mM HEPES. Prior to imaging, CAD cells were plated on coverslips coated with 10 μg/mL Laminin (Sigma).

### Immunofluorescence

Cells were fixed with 4% electron microscopy grade paraformaldehyde (PFA, Electron Microscopy Sciences, Hatfield, PA) for 10 min at RT, permeabilized with 0.2% Triton X-100 (Sigma-Aldrich) for 3 min, and washed twice with 1X DPBS. Cells were stained overnight at 4°C with primary antibodies diluted in immunofluorescence staining buffer. They were then washed twice with DPBS for 5 min, incubated with secondary antibodies (diluted 1:1000) for 1 hr at room temperature in immunofluorescence staining buffer. F-actin was stained with Phalloidin-568 (diluted 1:100, Life Technologies) for 30 min at room temperature in immunofluorescence staining buffer. Finally, cells were washed three times with DPBS before mounting with Prolong Diamond W/ DAPI (Life Technologies). We used the following antibodies/stains: Mouse anti-β3 Tubulin (TUJ1 1:500 dilution, Covance, Princeton, NJ), Goat anti-ChAT (1:1000, AB144, ED Millipore), Rabbit anti-Islet1 (1:1000, NBP2-14999, Novus Biologicals), Mouse anti-Tau (1:25000, gifted from the Giasson lab (Strang et al., 2017)) Alexa Fluor™ Phalloidin 568, anti-mouse IgG 488 and anti-rabbit IgG 488 (Life Technologies) used at 1:1000.

### Microscopy

High resolution images of motor neurons and CAD cells were acquired with a Nikon A1R+ laser scanning confocal microscope with 40X 1.3 NA or 60X 1.49 NA objectives and a GaAsP multi-detector unit. Imaging of cells for outgrowth and branching pattern analysis was done using the EVOS XL digital inverted microscope (Life Technologies).

### Image analysis

#### Neurite growth cone analysis

Confocal z-stacks were converted into a single maximum intensity projection image. Terminal neurite growth cone size, filopodia length, and filopodia number were analyzed using Fiji (ImageJ) software. Filopodia length was defined as the distance between lamellipodium edge to the furthest end of the extending filopodia. Values were exported into Microsoft Excel and GraphPad Prism for statistical analysis.

#### Neurite tracing and branching analysis

Images were taken on the EVOS XL digital inverted microscope and imported into Fiji (ImageJ) software. All visible projections in these images were traced using the Simple Neurite Tracer plugin (Longair et al., 2011). An image stack was created from the tracing which was then analyzed using the Sholl analysis plugin (Ferreira et al., 2014). To compare the relative change in neuron radius between NTg and SOD1 mutants across different experiments each set was normalized to the average radius of the age matched NTg control group. The Sholl profile containing the number of branches per given distance from soma and overall neuron radius was then exported to Microsoft Excel or GraphPad Prism for analysis.

#### Axon identification and analysis

Staining of motor neuron cultures were performed as previously described with a mouse-anti tau antibody to specifically identify the axon. Imaging was performed on the EVOS XL digital inverted microscope and the images were imported into Fiji (ImageJ) software. The 16-bit color lookup table was applied to each image to visualize the intensity of tau-staining in the neurites. The axon (neurite that was most intensely stained for tau) was then measured from the center of the soma to determine if it was the longest projection from the cell. We found that after two days in culture the process furthest from the soma correlated with highest tau expression 68% of the time (Supplementary Figure 2), leading us to conclude that in the majority of our analysis, overall cell radius directly correlated to axonal outgrowth.

### Adeno-associated virus (AAV) mediated overexpression of mutant SOD1

Motor neurons from adult (9-12 months) NTg mice were isolated and plated as described above. On the day they were plated, 20 μl of AAV2/8 (titer 1X10^9^) expressing wild type SOD1-YFP, SOD1^G93A^-YFP and GFP only was added to the wells and the cells were cultured for 10 days (growth medium was refreshed every 5 days). For these experiments we used a self-complementary virus (scAAV), driven by the chick beta actin promotor (CBA) as described in (Rosario et al., 2016). After 10 days in culture, cells were fixed and stained for β3 Tubulin then imaged/analyzed as described above.

### Cath.-a-differentiated (CAD) cell filopodia analysis

CAD cells were transfected 12-24 hrs prior to imaging with the appropriate constructs using the Neon electroporation system (Invitrogen) using a single 1400 v 20 ms pulse. 1 μg of DNA was used for each 10 μL electroporation. This protocol routinely gave >99% transfection efficiency and <10% cell death. YFP, wild type human SOD1-YFP, and human SOD1^G93A^-YFP constructs were expressed from plasmids based on the pEF-BOS expression vector. After transfection, CAD cells were cultured in serum-free medium for 18 hours to induce the formation of neurite-like processes. Cells were then fixed and stained for phalloidin as described above. Neurite-like processes were imaged using Nikon A1R+ laser scanning confocal microscope with 60X objective. The phalloidin channel of each image was deconvolved using Nikon Elements to enhance the resolution of filopodia selection. The ImageJ plugin Filopodyan (Urbančič et al., 2017) was then used to individually select filopodia and measure their YFP fluorescence.

## ACKNOWLEDGMENTS

This project was supported by a grant from the National Institutes of Health (NIH) (R01 NS092788) (D.R.B), a NIH Pathway to Independence Award (R00 NS087104) (E.A.V.) and a Starter Grant from the ALS Association (E.A.V and T.A.R)

